# Explosive diversification following continental colonizations by canids

**DOI:** 10.1101/2021.01.08.425986

**Authors:** Lucas M. V. Porto, Rampal S. Etienne, Renan Maestri

**Affiliations:** Groningen Institute for Evolutionary Life Sciences, University of Groningen, PO Box 11103, 9700 CC Groningen, The Netherlands; Ecology Department, Universidade Federal do Rio Grande do Sul, PO Box: 15007, Porto Alegre, Rio Grande do Sul, Brazil

**Keywords:** Ancestral range estimation, Canidae, Dispersion routes, Ecological opportunity, Evolutionary radiation, Geographical distribution

## Abstract

Colonization of a new environment may trigger an explosive radiation process, defined as an accelerated accumulation of species in a short period of time. However, how often colonization events trigger explosive radiations is still an open question. We studied the worldwide dispersal of the subfamily Caninae, to investigate whether the invasion of new continents resulted in explosive radiations. We used a combination of phylogenetic analyses and ancestral area reconstructions to estimate ancestral ranges of 56 extant and extinct species of Caninae, as well as variation in speciation and extinction rates through time and across clades. Our findings indicate that canids experienced an explosive radiation event when lineages were able to cross the Bering Strait and the Isthmus of Panama to reach Eurasia and South America, respectively, around 11 million years ago. This large number of species arising in a short period of time suggests that canids experienced ecological opportunity events within the new areas, implying that the differences in the ecological settings between continents may be responsible for the variation in clade dynamics. We argue that interaction with other carnivores probably also affected the diversification dynamics of canids.

## 1 Introduction

Evolutionary radiations are phenomena in which a large number of species arise in a short period of time (Lovette and Bermingham 1999, Schluter 2000, Rabosky and Lovette 2008, Losos 2011). Such radiations have contributed substantially to the earth’s biodiversity (Lovette and Bermingham 1999; Rabosky and Lovette 2008; Losos 2011; Morlon et al. 2012). Evolutionary radiations are often associated with the colonization of initially empty ecological space as a result of ecological opportunity, i.e. when species have access to new resources that are little used by other taxa (Schluter 2000). Geographic colonization of areas previously unoccupied by potential competitors or occupied by competitively inferior organisms is therefore an ideal setting for an evolutionary radiation to unfold (Stroud and Losos 2016), i.e. to generate cladogenesis accompanied by ecological and morphological disparity among lineages (Harmon et al. 2003).

Detecting evolutionary radiations requires knowledge of how speciation and extinction rates varied over time (Rabosky and Lovette 2008). The common assumption is that diversification is highest at the beginning of the radiation (Schluter, 2000; Etienne & Haegeman, 2012). In the last decades, the number of robust and reliable molecular phylogenies has increased together with methods to extract information about the tempo and mode of evolution from them (Nee 2006; Rabosky et al. 2007; Etienne and Haegeman 2012; Etienne and Rosindell 2012; Etienne et al. 2012; Revell 2012; Yu et al. 2015; Morlon et al. 2016; Herrera-Alsina et al. 2019). Patterns of evolutionary radiations have been identified for a variety of organisms (Burbrink and Pyron 2010; Wagner et al. 2012; Tran 2014; Arbour and López-Fernández 2016; Maestri et al. 2017; van Els et al. 2019). Once the timing of shifts in diversification rates has been established, one can compare them to major shifts in biogeographical distribution that characterize entrance in new geographic areas to find support for the hypothesis that this entrance triggered the radiation.

An ideal group to investigate patterns of diversification in continental evolutionary radiations is that of the Canidae, because their evolutionary history is marked by episodes of dispersal and colonization of new environments. Furthermore, canids are distributed all over the planet, have a rich fossil history, and well-resolved phylogenetic relationships between both extinct and extant species (Finarelli 2007; Porto et al. 2019). The family Canidae originated in North America approximately 40 million years ago (Mya), an epoch where this continent did not have land connections to any other continent (Wang and Tedford 2008; Prothero 2013). Within Canidae, successive radiations gave rise to three subfamilies by the end of the Oligocene: Hesperocyoninae, Borophaginae, and Caninae. The first two are now extinct and were restricted to North America, while Caninae radiated over almost the entire planet allowed by two geological events (Geffen et al. 1996; Cox 2000; Sillero-Zubiri et al. 2004; Wang and Tedford 2008; Potter and Szatmari 2009): the emergence of the Bering Strait and the Isthmus of Panama, both around 11 Mya (MacNeil 1965; Hopkins 1967; Montes et al. 2015). Today, the Caninae is separated into three major clades: true-wolves, Cerdocyonina, and Vulpini (which we hereafter will refer to as wolves, South American canids, and foxes, respectively) (Wang and Tedford 2008).

In this study, we used phylogenetic information of the subfamily Caninae to investigate the timing and location of major dispersal events, to explore the biogeographical processes that led to the present canid distribution, and to associate biogeographical events with estimates of shifts in speciation and extinction rates through time and across clades.

## 2 Materials and Methods

### 2.1 Taxon sampling and phylogenetic information

We used the phylogenetic tree from Porto et al. (2019) as the base for our study. This tree was constructed with molecular and osteological data, through Bayesian inference, and presents all the 37 extant canids in the world. To increase accuracy in the ancestral range estimation, we added 19 extinct species (Table S1) to this tree following taxonomic information from the digital paleobiology dataset, *Fossilworks* (Alroy 1998). We performed a literature review in *Fossilworks* in search of articles that presented information on the most likely taxonomic position that the extinct species would fit into. With this information we added the species to the phylogeny at the most likely nodes, resulting in a phylogeny of 56 species (representing 46.7% of all known species of the Caninae subfamily, based on Wang & Tedford (2008)). We only added extinct species that had a detailed description of their past geographical distribution in the literature, and thus we could generate maps (polygons) with the occurrence points to classify these species into the eight biogeographic regions used here (Fig. 1). We dated the phylogeny again, using the fossil information of the 19 species and the fossil ages that were used by Porto et al. (2019). We used *Leptocyon vafer* (Leidy 1858) and *Leptocyon vulpinus* (Matthew 1907) as the outgroup for the biogeographical analyses because it is known that both species originated in North America during the Miocene, so the root of the phylogeny will be fixed where the subfamily Caninae originated. The phylogeny is shown in Fig. S1.

**Figure 1.**
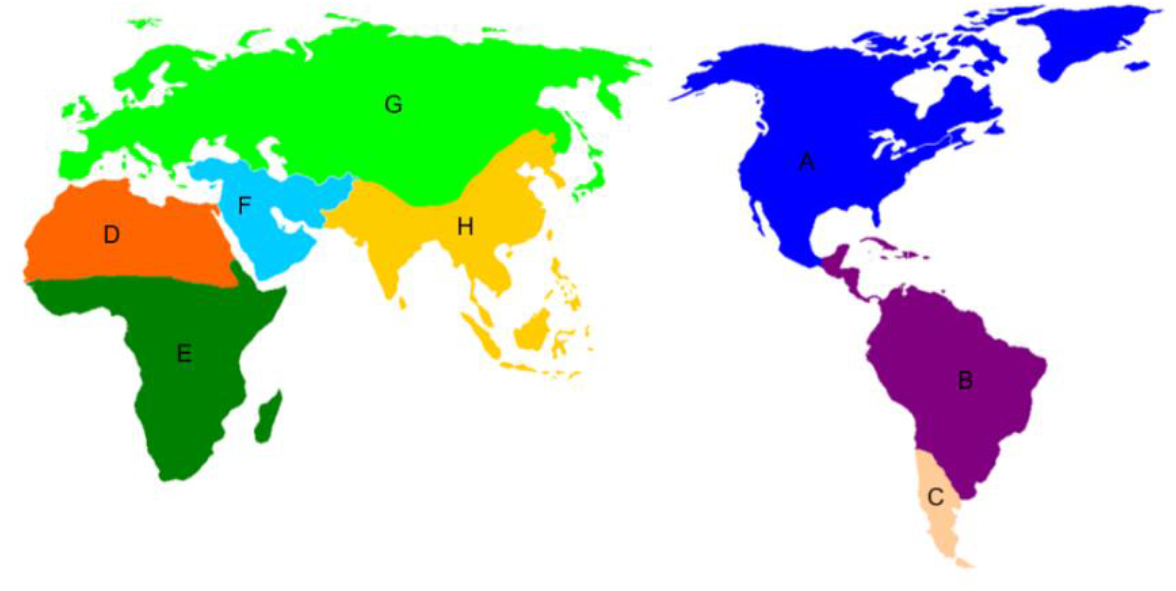
Eight biogeographic regions delimitated for our ancestral range estimation analysis based on the patterns of the beta-sim index of mammal species from Kreft & Jetz (2010), and the distribution shapes from IUCN (2017). The regions delimitated are North America (A), Northern South America (B), Southern South America (C), Northern Africa (D), Sub-Saharan Africa (E), Middle East (F), Northern Eurasia (G), and Southern Eurasia (H).

### 2.2 Ancestral range estimation

To estimate ancestral ranges, we divided the world into eight biogeographic regions (Fig. 1) based on the patterns of the beta-sim index of mammal species from Kreft & Jetz (2010), and on the distribution shapes of the extant species taken from IUCN (2008). We categorized all the 56 canids into one or more biogeographical regions based on their distributions as given in (IUCN 2008) (Table S1).

We performed an ancestral range estimation using *RASP* 4.0 (Yu et al. 2015). Six models of ancestral range estimation were fitted and compared: The Dispersal-Extinction Cladogenesis Model (DEC), a likelihood version of the Dispersal-Vicariance Analysis (DIVALIKE), the likelihood version of the Bayesian Analysis of Biogeography (BAYAREALIKE), as well as “+J” versions of these three models, that include founder events of speciation. DEC assumes that one daughter lineage will always have a range of one area and, at least, one daughter lineage inherits the range of the ancestor lineage. DIVALIKE allows that all daughter lineages have two or more areas, but this model does not allow sympatric speciation. BAYAREALIKE assumes the daughter lineages will have the same range as their ancestor. For more details on the assumptions of these different models see Matzke (2013). We compared the predictions of these different models rather than selecting the best-fit model based on the Akaike Information Criterion corrected for small sample sizes (AICc) (Akaike 1973) because it has been argued that this model selection may be biased (Ree and Sanmartín 2018). The number of sampled trees was set to 1 × 10^6^.

To understand how canids dispersed worldwide, we used the informative matrices (Appendices S1 and S2) of the ancestral range estimation analysis, which summarize the most likely paths by which ancestral lineages followed. These matrices contain all the events that occurred in each ancestral node of the tree (Yu et al. 2015). Based on this data, we derived, in a form of a palaeoscenario, the dispersal routes that the lineages of Caninae followed during their evolutionary history.

### 2.3 Speciation and extinction rates

We estimated speciation and extinction rates through time and across clades using the Bayesian Analysis of Macroevolutionary Mixtures version 2.6 (*FossilBAMM*) (Mitchell et al. 2019). *FossilBAMM* implements an automatic reversible-jump Markov Chain Monte Carlo algorithm (rjMCMC), which enables the detection of changes in the rates of speciation and extinction of canids, allowing us to visualize rate peaks over time with non-ultrametric trees, even with an incomplete fossilized history. Speciation and extinction rates were calculated for the whole tree and separately for the three major clades of Caninae (wolves, foxes, and South American canids). Each analysis was performed using four chains, with 20 × 10^6^ iterations, and samples were obtained every 1000 cycles. We removed the first 20% of the collected samples as burn-in. We used the *BAMMtools* package (Rabosky et al. 2014) in the *R* environment (R Development Core Team 2020) to plot and visualize the results from *BAMM*. that allows estimates to be made. To detect rate shifts over the tree, we extracted the potential rate shifts and associated parameters, together with their relative probabilities, from all the collected samples (80%).

While *BAMM* has been criticized in the literature (Moore et al., 2016), Rabosky et al., (2017) have argued that previous criticisms about *BAMM* are incorrect or unjustified, although it is true that the method, like many others in the same family of models, is not exactly correct (Laudanno et al. 2020). More importantly, over the last few years, several studies have demonstrated that most inferences seem to be robust when estimating diversification in distinct groups of animals (Rabosky et al. 2013; Shi and Rabosky 2015; Chang et al. 2019; Rabosky 2020).

## 3 Results

### 3.1 Ancestral range estimation

We provide a detailed analysis of the predictions of the DEC + J model and then compare the predictions of the other models to this model (Fig. 2B) because DEC + J was the model that presented less uncertainty during node reconstruction compared to the others. According to DEC + J, the ancestor of the three major clades of Caninae originated in North America, but the first lineage to disperse out of this continent was the lineage that led to the South American canids, around 10.4 Mya. After a long period, diversifying in the north part of South America, the South American canids dispersed to Patagonia through three lineages (Fig. 3A - M2, M4, and M6), around 4 and 3 Mya, originating four species. Two South American lineages dispersed back to North America around 4.8 Mya (Fig. 3A - M3 and M5), giving rise to *Chrysocyon nearcticus* and *Cerdocyon avius*.

**Figure 2.**
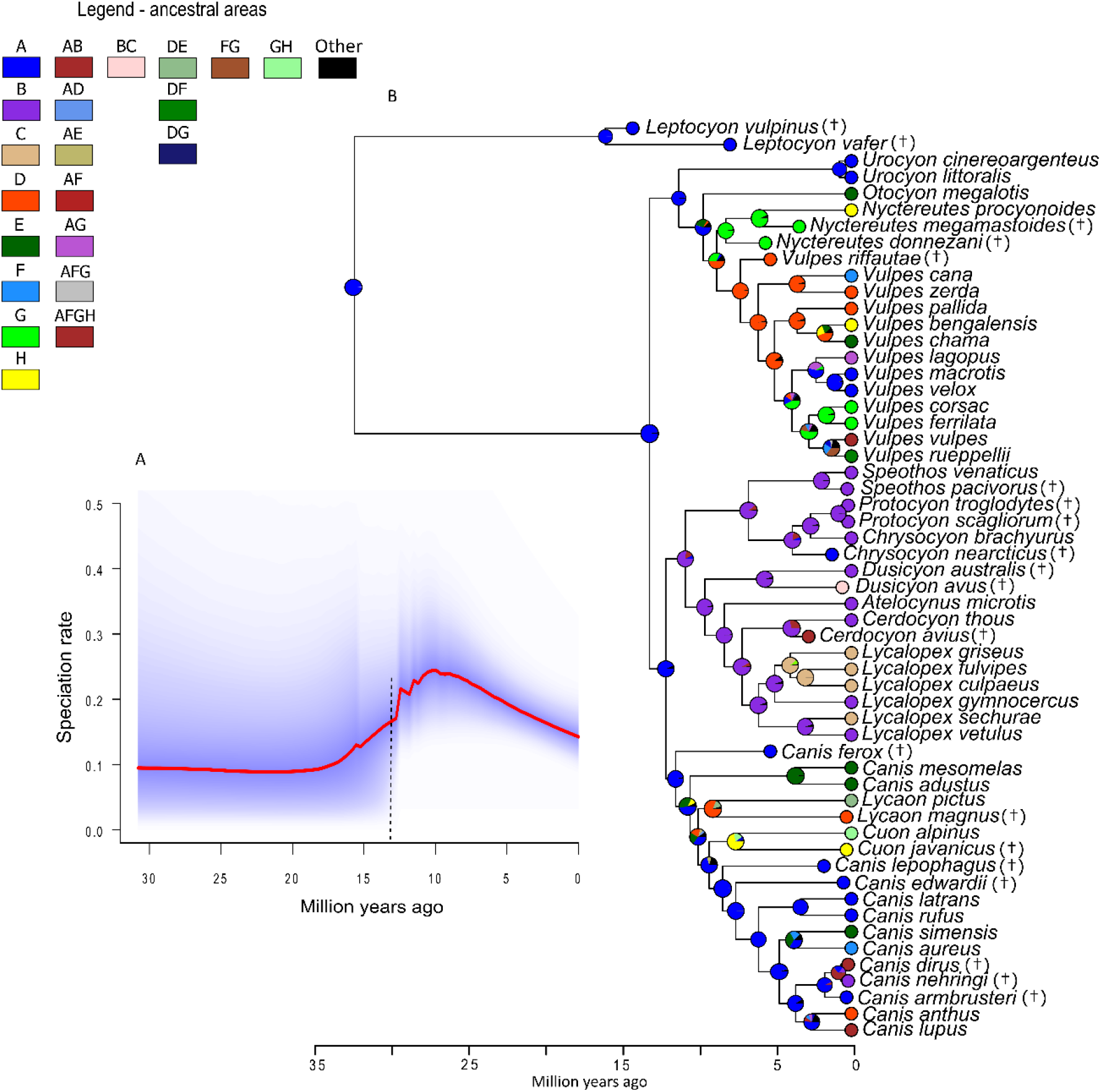
Speciation rate for the whole tree (A), phylogenetic tree, and ancestral range reconstruction under DEC + J (B). The probability of the ancestral areas of the lineages is indicated at the nodes of the tree, and the color-coded circles at the tips represent the current areas occupied by each lineage. The colors represent the different biogeographic regions as indicated in the legend (left). The black dashed line indicates the time of significant rate shift (A). The species with the symbol (†) are the extinct species added to the tree.

**Figure 3.**
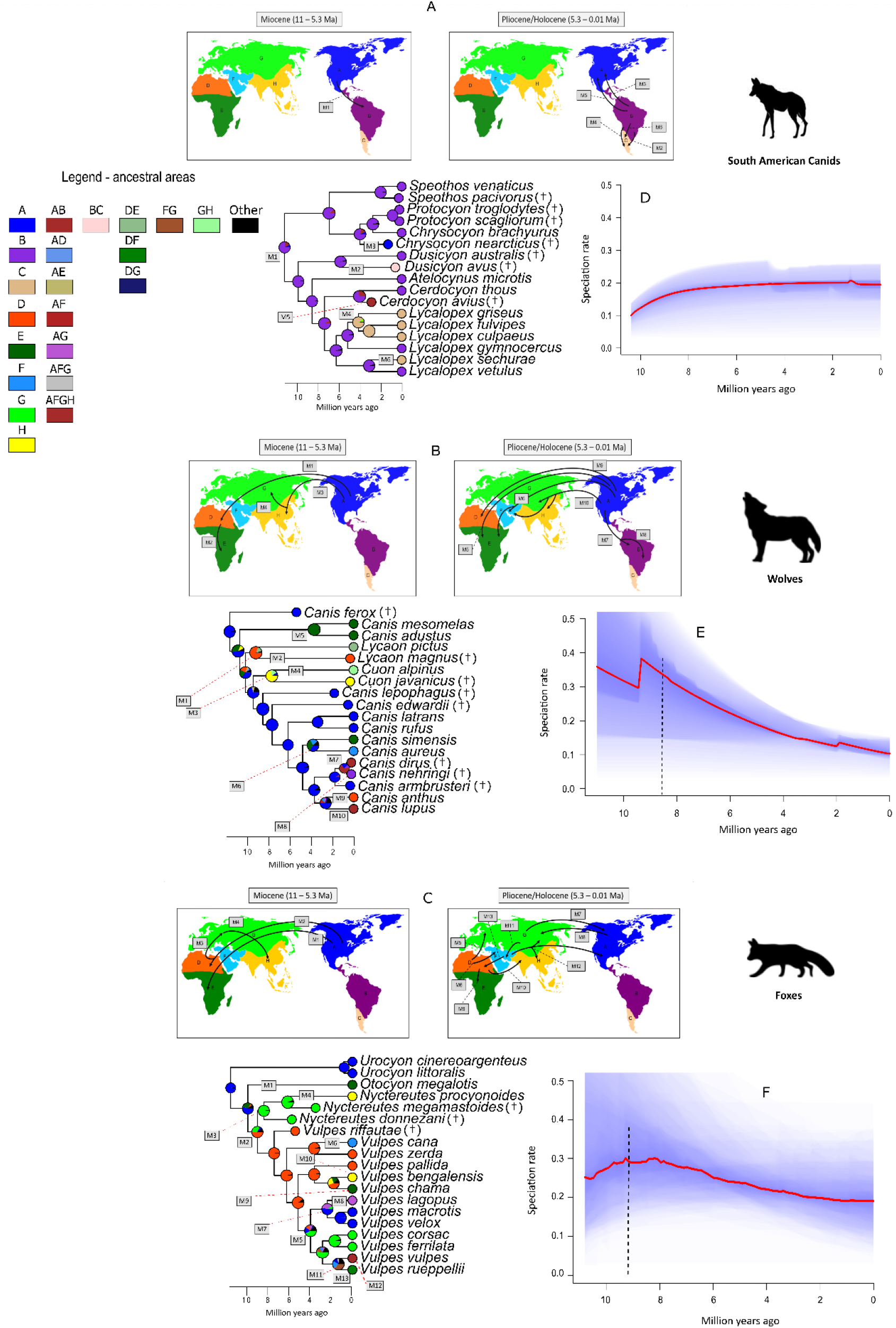
Ancestral range estimation for the South American canids (Cerdocyonina) (A), wolves (B), and foxes (C) under DEC + J model. The hypothetical dispersal routes that lineages used are indicated by black arrows. The probability of the ancestral areas of the lineages is indicated at the nodes of the tree, and the color-coded circles at the tips represent the current areas occupied by each lineage. The respective areas that the colors represent are indicated in the maps. All the dispersal events that occurred over the tree are coded as moments (M1, M2…M*x*) and are also indicated in the maps at the respective times that occurred (Miocene or Pliocene/Holocene). We also show the speciation rates for the three major clades: South American canids (D), wolves (E), and foxes (F). The peaks in the speciation rates occurred ~ 9 Mya for the three major clades separately. Black dashed lines indicate the time of significant rate shifts (D, E, and F). The species with the symbol (†) are the extinct species added to the tree.

Wolves went for the first time to the Old world through two lineages between 9.5 and 7 Mya, which dispersed to Africa and Asia, respectively (Fig. 3B - M1 and M3), and gave rise to four species that were distributed in a great part of these continents. Wolves experienced a long period diversifying within North America until around 4.5 Mya, when a lineage (Fig. 3B - M5) dispersed to Sub-Saharan Africa. *Canis lupus* and *Vulpes vulpes*, which are the species with the largest current biogeographic distributions, originated from endemic lineages of North America, dispersing to other areas in a relatively short time, 3.4 Mya. Within the wolf clade, the extinct species of *Canis dirus* and *Canis nehringi* originated from a lineage that dispersed to South America around 1 Mya (Fig. 3B - M7 and M8).

Foxes also originated in North America, but dispersed to the Sub-Saharan Africa around 9 Mya (Fig. 3C - M1). The most important dispersal event for foxes was the event M2, around 8.5 Mya when a lineage went to North Africa. From this ancestor, the foxes started a diversification process inside North Africa generating new lineages in this area, but also dispersing to other areas, such as Eurasia (Fig. 3C - M3 and M5) and North America (Fig. 3C - M7). Unlike the wolves, foxes had their center of diversification in North Africa.

The other five biogeographic models used here estimated ancestral ranges in slightly distinct ways. DIVALIKE + J resembles the DEC + J generating a similar ancestral estimation for foxes and South American canids. However, under DIVALIKE + J, the ancestral lineages of wolves had an area expansion, around 11 Mya, inhabiting a great part of the north-hemisphere. Then, around 9 Mya, it seems that a retraction in their distributions occurred, originating more geographically restricted lineages (Fig. S2). BAYAREALIKE + J is similar to DEC + J for the diversification of both wolves and foxes (Fig. S3). They differ in predictions for the South American canids. The estimation of BAYAREALIKE + J suggests that, for this clade, the first lineages inhabited a wide distribution (North America and a great part of South America), different from the DEC + J model that shows the diversification of this clade occurring within South America only. The DEC, BAYAREALIKE, and DIVALIKE models (Figs. S4, S5, and S6), in which the J parameter is not present, are all similar to DEC + J for the clade of foxes and South American canids. However, they all differ from the DEC + J model because the predictions for wolves were more similar to those of DIVALIKE + J, where the ancestral lineages of wolves had an expansion and then a retraction in their distributions before originating the extant lineages.

### 3.2 Speciation and extinction rates

Peaks in speciation rates through time were coincident for Canids (whole tree) and each of its major subclades (Figs. 2A, 3D, 3E, and 3F), around 12 - 8 Mya. Therefore, changes in speciation rates were triggered in parallel across different lineages. The peak for the Caninae tree starts at 12 Mya (λ = 0.25). The results for the three clades separately showed that in wolves, foxes, and South American canids the peaks occurred at ~ 9 Mya (λ = 0.37; λ = 0.28 and λ = 0.16). After the rate peaks, South American canids stabilized their speciation rate, whereas the speciation rate of wolves and foxes declined. For the whole tree, the extinction rate remained at its highest value until 17 Mya (μ = 0,13), and after that, the rate began to decline over time (Fig. S7A). The extinction in wolves presented a peak 6 Mya (μ = 0,12), while in foxes and South American canids the extinction rates of both clades were constant over time (μ = 0.1 and μ = 0.02) (Fig. S7B, S7C and S7D).

We detected three significant rate shifts during the speciation of Canidae: one for the whole tree (Fig. 2A) and the other two for the clades of wolves and foxes, respectively (Figs. 3E and 3F). The uncertainty during the speciation rate estimations was higher when clades were analyzed separately due to the removal of many species for the comparison between clades.

## 4 Discussion

Our findings show that major events of canid dispersal — to South America and North Africa — are associated with peaks in diversification rates, suggesting an evolutionary radiation process right after the geographic colonization of new continents. Moreover, the pattern presented by the speciation rates in the clades of wolves and foxes, as well as for the whole tree, with a decline in the speciation rate after a peak in the emergence of new lineages is characteristic of an explosive radiation (Schluter 2000; Rabosky and Lovette 2008). Our results contrast with those of Liow & Finarelli (2014), which showed stable diversification rates of carnivores over the last 22 Ma.

The speciation rate of Caninae increased substantially shortly after canids arrived in the Old World and South America around 10 Mya (Fig. 2A). The timing of radiations suggests that this pattern for the whole tree was generated by ecological opportunity after the entrance in new continents by South American canids and foxes, but also by the diversification of wolves within North America. For the first two clades, the peaks in speciation rates occurred at the same time as these lineages entered in North Africa and South America (Fig. 3D and 3F). The absence of competitors and the new types of vegetation in Africa (deserts) (Zhang et al. 2014) and South America (tropical forests), may have generated the ecological opportunity and subsequent Caninae diversification in these continents. This ecological opportunity hypothesis must still be tested with ecological data.

Wolves had their peak around 9 Mya, but it seems that their explosive diversification was not triggered by the entrance in new areas, but probably due to the great turnover in the North American herbivorous fauna associated with the expansion of grasslands, resulting in ecological innovations in canids (Cox 2000; Janis et al. 2000; Figueirido et al. 2015). As our results showed, during the Miocene, there were not many dispersion events among biogeography areas that could explain the peak in wolves` speciation (Fig. 3B). Therefore, the explosive diversification that wolves underwent in North America was probably due to the changing environment in this continent rather than the colonization of new areas.

We note that not all dispersal events to new areas led to radiations. Dispersal events M3 and M4 in wolves and M4 in foxes, all in Eurasia, did not increase the speciation rates of both clades (Figs. 3B, 3C, 3E, and 3F). It is likely that the large number of other carnivores in this area (e.g., Felidae, Barbourofelidae, and Hyaenidae) (Werdelin and Solounias 1991; Zhanxiang 2003; Wang and Tedford 2008) imposed strong competition on canids, which may have resulted in a slowdown in the rate of speciation of these clades. Intense competition with other carnivores may explain why Eurasia was not the center of diversification for any of the three major clades of Canidae.

In addition, wolves and foxes showed a decrease in the speciation rate, as well as the whole tree, after their peak (Fig. 2A, 3E, and 3F). This brings us to another characteristic of explosive radiations, declining speciation through time due to a presumed saturation of available ecological opportunity (Schluter, 2000; Harmon et al., 2003; Rabosky & Lovette, 2008, Etienne et al. 2012). In foxes, a plausible explanation would be that niches became filled over time, mainly in North Africa (diversity-dependence). Wolves, mainly in North America, may have experienced competition with other carnivores that came from Eurasia, such as felids (Johnson et al. 2006). These interactions may have contributed to the decay in the speciation rate and the increase in the extinction rate of wolves (Silvestro et al. 2015; Pires et al. 2017), as biotic interactions such as competition can constrain evolutionary dynamics (Pires et al. 2015). Wolves in North America may also have had their dispersal out of the continent limited by the carnivores in Eurasia (Werdelin and Solounias 1991; Zhanxiang 2003; Wang and Tedford 2008) through an incumbency effect (Rosenzweig and Mccord 1991), which could explain the great diversification of wolves only in North America.

The three major clades of canids showed distinct dispersal patterns over the world. Wolves and foxes, even though they have very similar geographic distributions nowadays, did not coexist in the same biogeographical regions for much time in the past. Because of the large number of speciation events in wolves on the North American continent, it is evident that the center of diversification of the wolves was North America, while foxes had their center of diversification in North Africa. Furthermore, our results suggest different arrival times of some species in certain biogeographic regions than previously known in the literature. For example, the ancestral range estimation indicated that the first canid to arrive in the Old World was the ancestor of both species, *Lycaon magnus* and *Lycaon pictus* (Fig. 3B - M1) about 9 to 8.5 Mya, differently from Crusafont-Pairó (1950) who proposed, with fossil data, that the *Canis cipio* as the first canid to disperse out of North America around 8 to 7 Ma.

Foxes and South American canids seem to have undergone a diversification process distinct from wolves. The diversification center of foxes, North Africa, is a desert environment that originated around 7 Mya (Zhang et al. 2014). Because it is a desert, food is probably very scarce for large predators such as lions and leopards, which now live in sub-Saharan Africa, but during the last five million years also inhabited parts of the Sahara (Johnson et al. 2006; Wilson and Mittermeier 2009; de Manuel et al. 2020). In this scenario, foxes may have had an ecological opportunity to occupy this area due to the generalist behavior that this clade evolved, as demonstrated by Porto et al. (2019), not overlapping their niches with other carnivores. A similar process probably occurred with the South American canids. Once in South America, the lineage that dispersed from North America faced an environment dominated by forest (Zachos et al. 2001; Potter and Szatmari 2009; Strömberg 2011) and a fauna composed mostly of large herbivores and lacking potential competitors (Wang and Tedford 2008).

Our results are based on separate analyses for the biogeographic ancestral reconstruction and for the shifts in diversification rates. Ideally, a single analysis should be used for this, such as GeoSSE (Goldberg et al. 2011) or other state-of-the-art SSE-type approaches (Beaulieu and O’Meara 2016; Herrera-Alsina et al. 2019), which can link the diversification rate directly to the biogeographic distribution of the lineage. However, these are currently computationally very demanding when there are many states which is the case here, and they have not been developed for phylogenies with fossil data. Similarly, the detection of ecological opportunity affecting diversification rates suggests the use of diversity-dependent diversification models (Etienne and Haegeman 2012; Etienne et al. 2012). This requires a combination of diversity-dependence diversification with ancestral state reconstruction, which has not yet been implemented.

## 5 Conclusion

We studied changes in speciation rates of Caninae in the light of distribution data to provide a detailed description of the dispersal and diversification of the subfamily Caninae through the world over the last 31 Ma. The spatial patterns indicated that Caninae underwent an evolutionary radiation process when entering in Africa and South America, suggesting that the differences in the ecological settings between continents may be responsible for the disparity among clades dynamics. We also suggest that the new environment arising in North America over the last 10 Mya was the major responsible for wolves` radiation rather than dispersion events outside this continent. Interaction with other carnivores, which came from Eurasia to North America, may have affected the speciation dynamics of wolves over the last 9 Mya.

## Acknowledgements

We thank Vanderlei Debastiani for the help with the speciation rate analyses. This study was financed in part by the Coordenação de Aperfeiçoamento de Pessoal de Nível Superior - Brazil (CAPES).

## DECLARATION SECTION

### Ethics approval and consent to participate

Not applicable.

### Consent for publication

Not applicable.

### Availability of data and material

All data generated or analyzed during this study are included in this published article and the supplementary information files.

### Authors’ contributions

L.M.V.P. conceived and designed the study and analyses. L.M.V.P. performed the analyses. L.M.V.P. wrote the first draft of the manuscript. R.S.E. and R.M. commented on the methods and contributed to substantial revisions on the draft.

