## Supporting information for "Explosive diversification following continental colonizations by canids"

### SUPPLEMENTARY TEXT: FIGURES

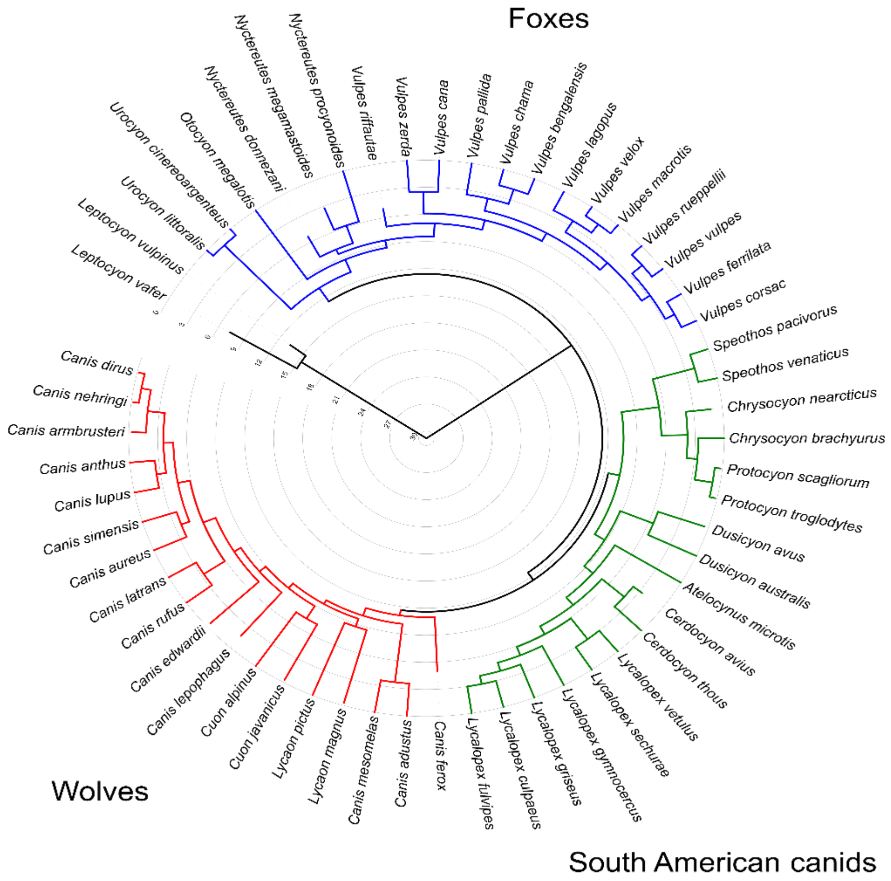


**Figure S1** Phylogenetic tree with 56 species of canids used during our analyses. This tree was constructed by adding 19 extinct species to the phylogeny of extant canids of Porto et al. (2019). The three major clades of Caninae are identified by the colors red (Wolves), blue (Foxes), and green (South American canids).

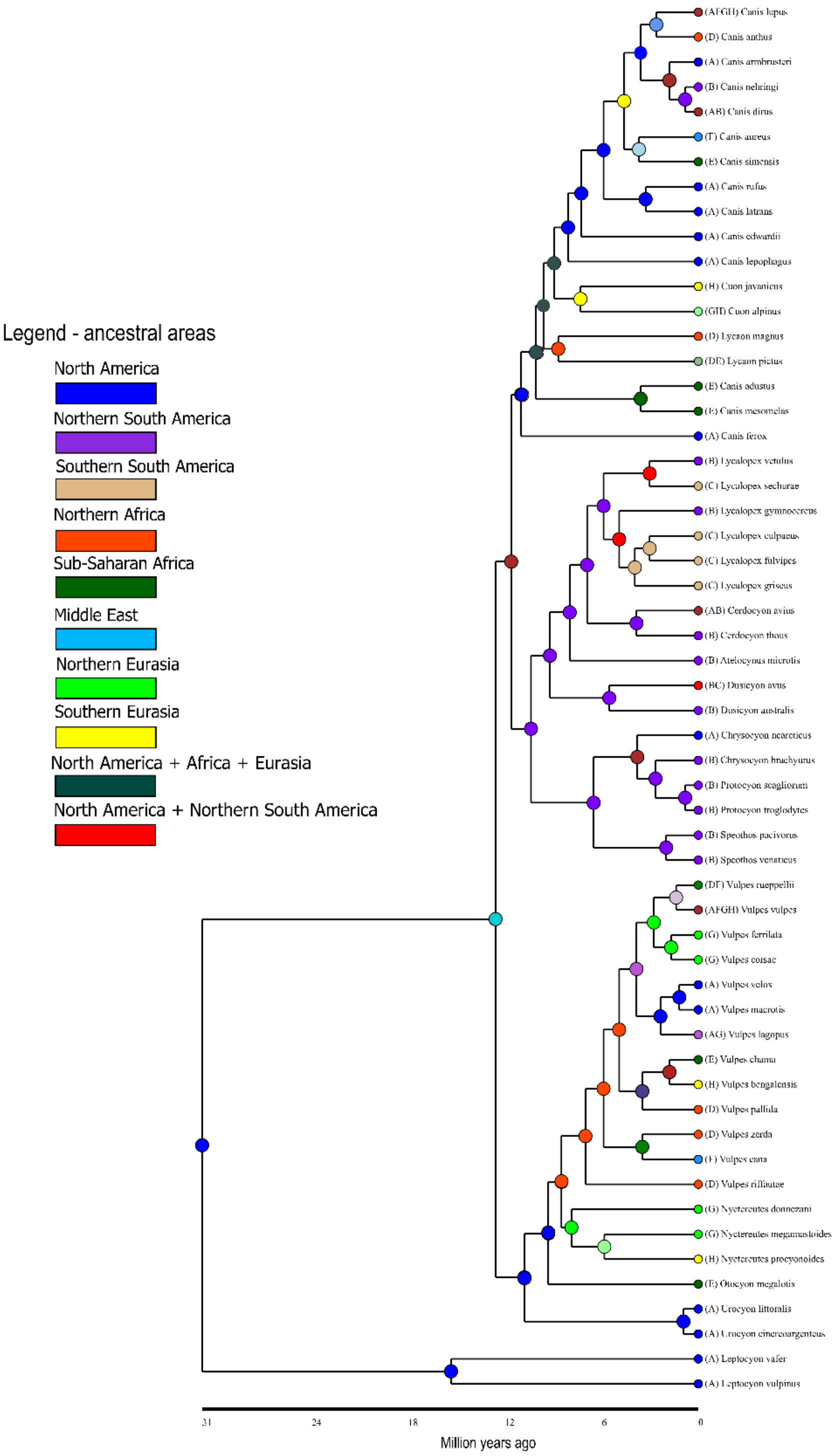


**Figure S2** Phylogenetic tree and ancestral range reconstruction for the whole tree under DIVALIKE + J. The ancestral areas of the lineages are indicated at the nodes of the tree and the color-coded circles at the tips represent the current areas occupied by each lineage. The colors represent the different biogeographic regions as indicated in the legend (left).

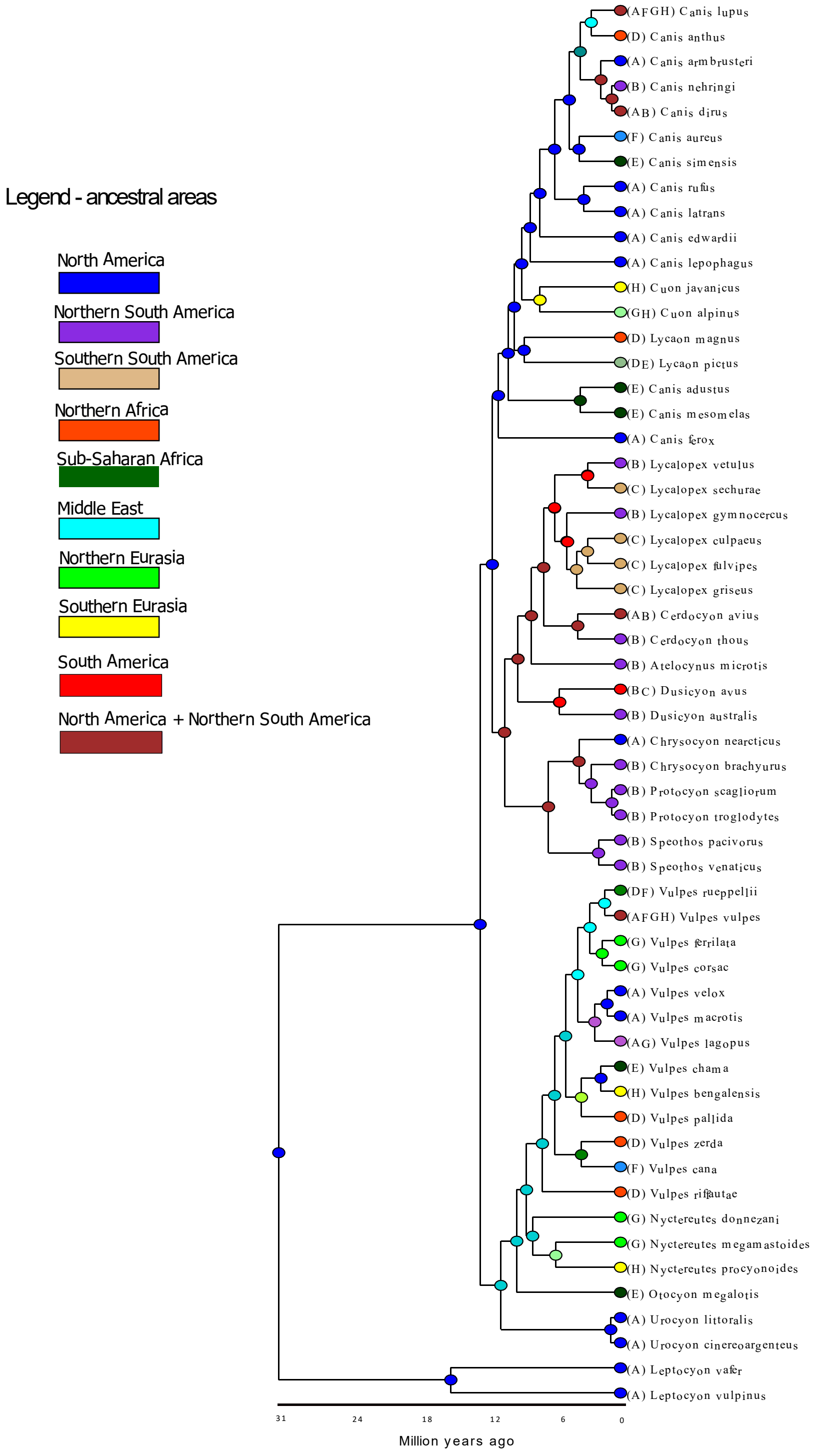


**Figure S3** Phylogenetic tree and ancestral range reconstruction for the whole tree under BAYAREALIKE + J. The ancestral areas of the lineages are indicated at the nodes of the tree and the color-coded circles at the tips represent the current areas occupied by each lineage. The colors represent the different biogeographic regions as indicated in the legend (left).

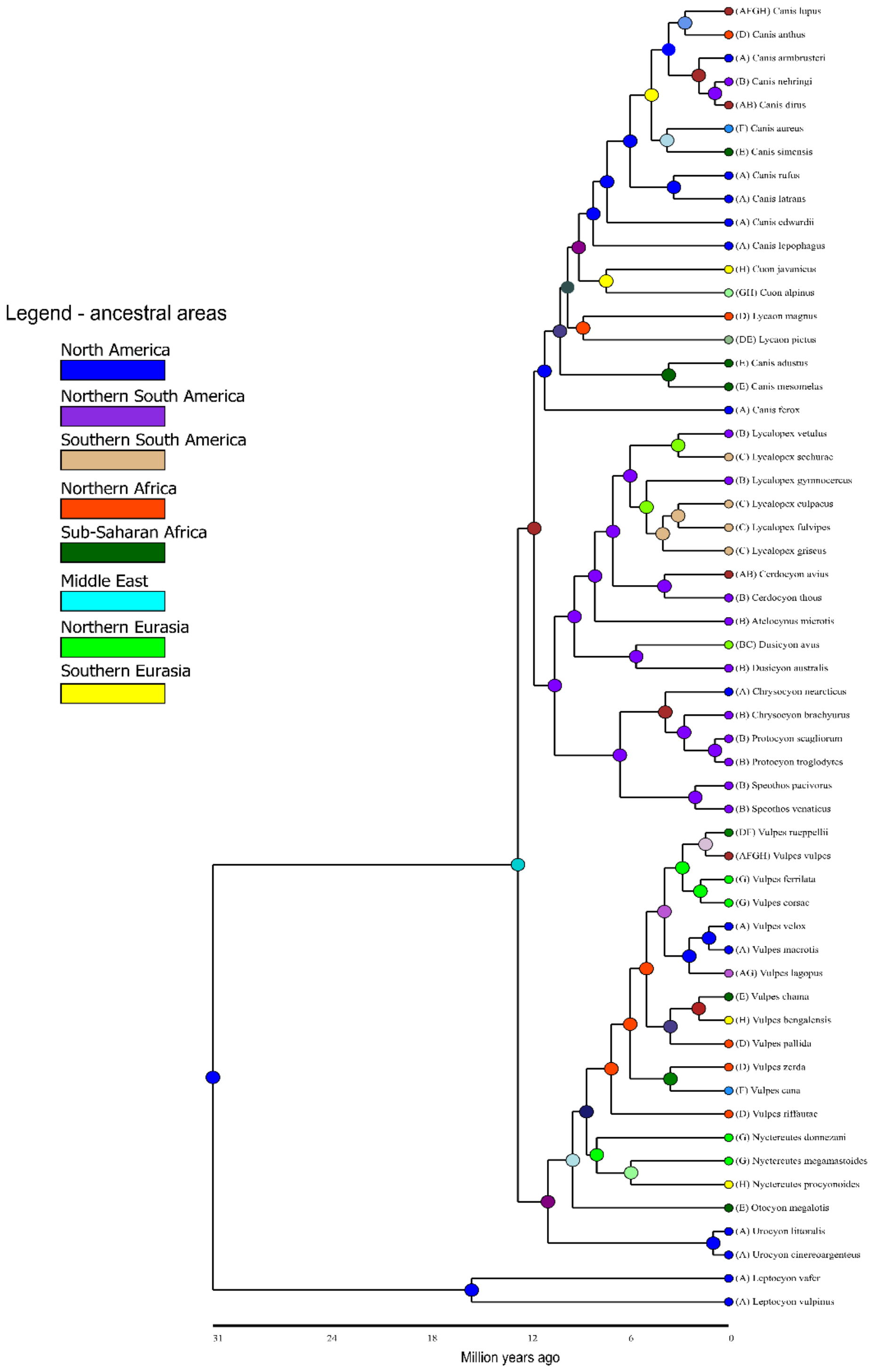


**Figure S4** Phylogenetic tree and ancestral range reconstruction for the whole tree under DIVALIKE. The ancestral areas of the lineages are indicated at the nodes of the tree and the color-coded circles at the tips represent the current areas occupied by each lineage. The colors represent the different biogeographic regions as indicated in the legend (left).

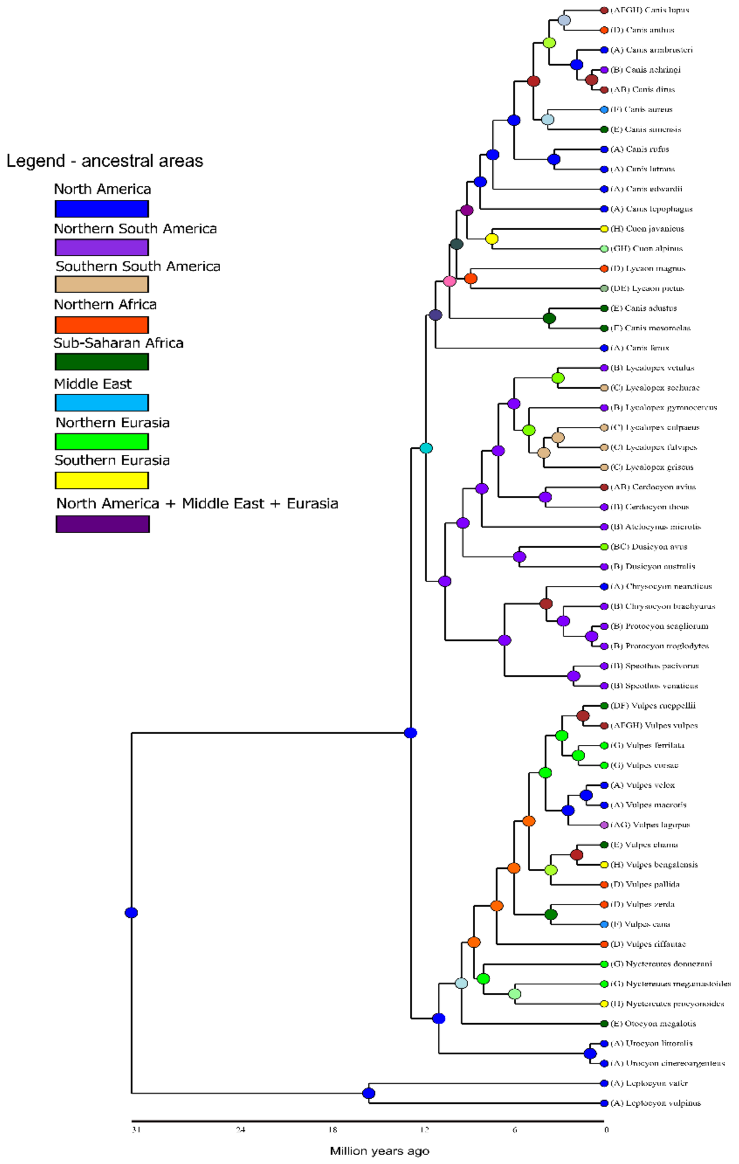


**Figure S5** Phylogenetic tree and ancestral range reconstruction for the whole tree under DEC. The ancestral areas of the lineages are indicated at the nodes of the tree and the color-coded circles at the tips represent the current areas occupied by each lineage. The colors represent the different biogeographic regions as indicated in the legend (left).

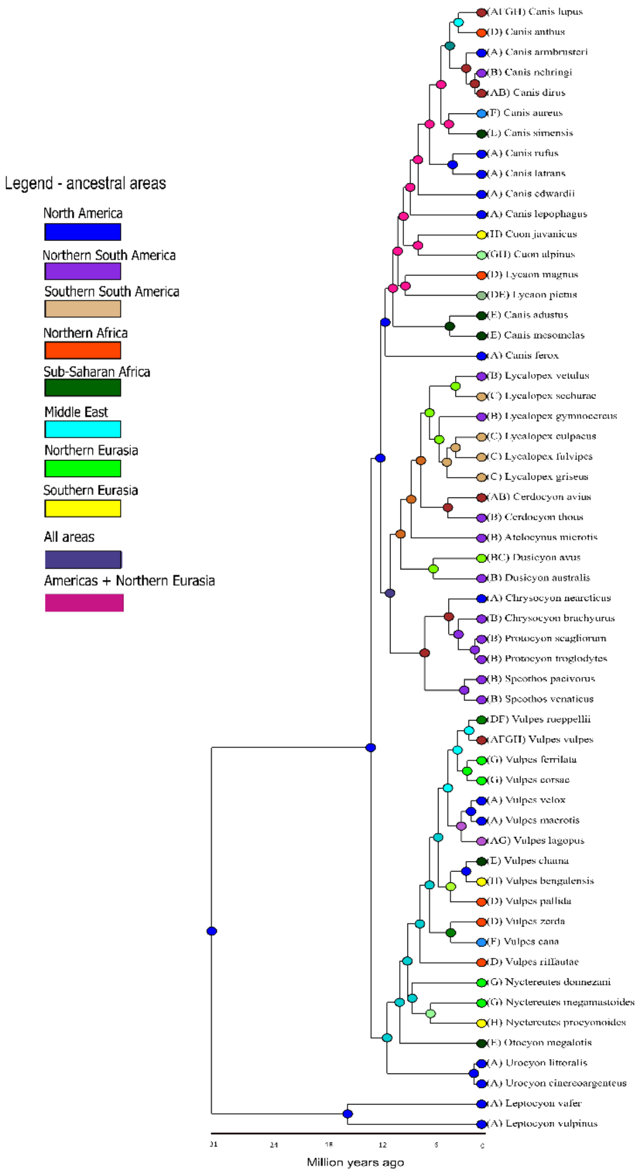


**Figure S6** Phylogenetic tree and ancestral range reconstruction for the whole tree under BAYAREALIKE. The ancestral areas of the lineages are indicated at the nodes of the tree and the color-coded circles at the tips represent the current areas occupied by each lineage. The colors represent the different biogeographic regions as indicated in the legend (left).

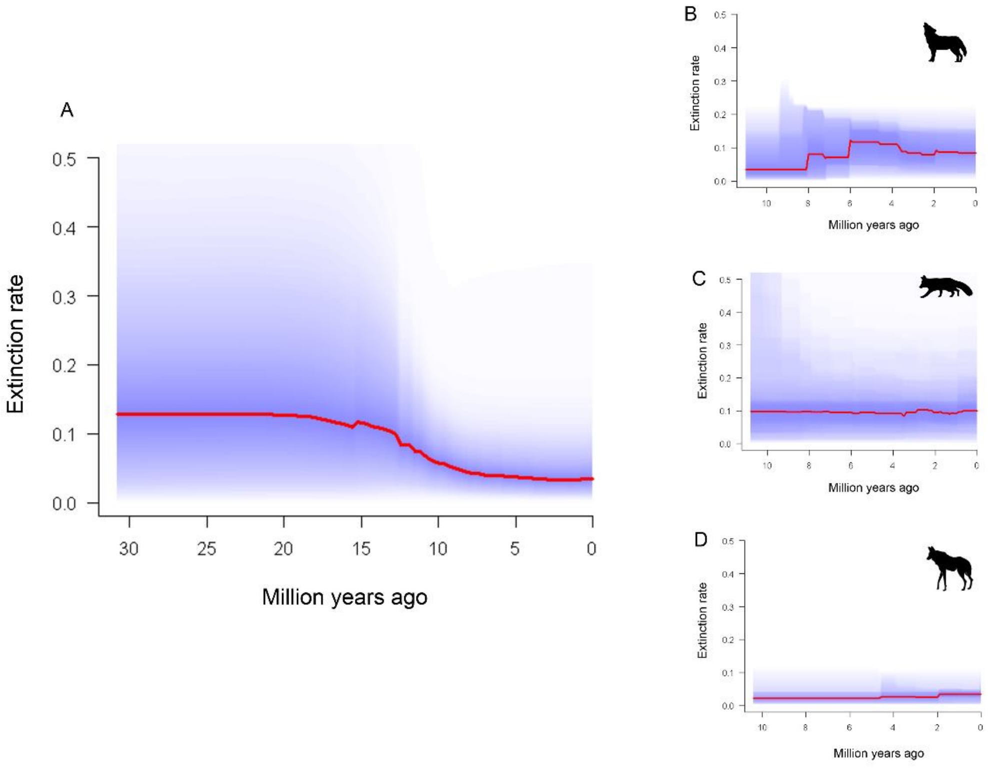


**Figure S7** Extinction rates for the whole tree (A) and for the three major clades of Caninae: wolves (B), foxes (C), and South American canids (D).

### SUPPLEMENTARY TEXT: TABLE LIST

**Table S1.** List of the 56 species of Canidae included in our study with the distribution areas that they belong based on our eight biogeographical regions (Fig. 1 in the main text). The original descriptor is also specified. Species marked with (*) are the 19 extinct canids include in the tree of Porto et al. (2019).

| Species | Biogeographic area | Descriptor |
|---|---|---|
| *Canis lupus* | ACE | Linnaeus, 1758 |
| *Canis anthus* | D | Cuvier, 1820 |
| *Canis aureus* | C | Linnaeus, 1758 |
| *Canis simensis* | D | Rüppell, 1840 |
| *Canis rufus* | A | Audubon and Bachman, 1851 |
| *Canis latrans* | A | Say, 1823 |

| | | |
|---|---|---|
| *Cuon alpinus* | E | Pallas, 1811 |
| *Lycaon pictus* | D | Temminck, 1820 |
| *Canis adustus* | D | Sundevall, 1847 |
| *Canis mesomelas* | D | Schreber, 1775 |
| *Lycalopex vetulus* | B | Lund, 1842 |
| *Lycalopex sechurae* | B | Thomas, 1900 |
| *Lycalopex gymnocercus* | B | Fischer, 1814 |
| *Lycalopex culpaeus* | B | Molina, 1782 |
| *Lycalopex fulvipes* | B | Martin, 1837 |
| *Lycalopex griseus* | B | Gray, 1837 |
| *Cerdocyon thous* | B | Linnaeus, 1766 |
| *Atelocynus microtis* | B | Sclater, 1883 |
| *Dusicyon australis* † | B | Kerr, 1792 |
| *Chrysocyon brachyurus* | B | Illiger, 1815 |
| *Speothos venaticus* | B | Lund, 1842 |
| *Vulpes rueppellii* | DC | Schinz, 1825 |
| *Vulpes vulpes* | ACE | Linnaeus, 1758 |
| *Vulpes ferrilata* | E | Hodgson, 1842 |
| *Vulpes corsac* | E | Linnaeus, 1768 |
| *Vulpes velox* | A | Say, 1823 |
| *Vulpes macrotis* | A | Merriam, 1888 |
| *Vulpes lagopus* | AE | Linnaeus, 1758 |
| *Vulpes chama* | D | Smith, 1833 |
| *Vulpes bengalensis* | E | Shaw, 1800 |
| *Vulpes pallida* | D | Cretzschmar, 1826 |

| | | |
|---|---|---|
| *Vulpes zerda* | D | Zimmermann, 1780 |
| *Vulpes cana* | C | Blanford, 1877 |
| *Nyctereutes procyonoides* | E | Gray, 1834 |
| *Otocyon megalotis* | D | Desmarest, 1822 |
| *Urocyon littoralis* | A | Baird, 1857 |
| *Urocyon cinereoargenteus* | A | Schreber, 1775 |
| **Canis dirus* † | AB | Leidy, 1858 |
| **Canis armbrusteri* † | A | Gidley, 1913 |
| **Leptocyon vafer* † | A | Leidy, 1858 |
| **Leptocyon vulpinus* † | A | Matthew, 1907 |
| **Cuon javanicus* † | E | Desmarest, 1820 |
| **Canis ferox* † | A | Miller and Carranza-Castaneda, 1998 |
| **Canis edwardii* † | A | Gazin, 1942 |
| **Lycaon magnus* † | D | Ewer and Singer, 1956 |
| **Canis lepophagus* † | A | Johnston, 1938 |
| **Vulpes riffautae* † | D | de Bonis et al., 2007 |
| **Cerdocyon avius* † | AB | Torres and Ferrusquia, 1981 |
| **Chrysocyon nearcticus* † | A | Tedford et al., 2009 |
| **Dusicyon avus* † | B | Burmeister, 1866 |
| **Canis nehringi* † | B | Ameghino, 1902 |
| **Protocyon troglodytes* † | B | Lund, 1838 |
| **Protocyon scagliorum* † | B | Giebel, 1855 |
| **Nyctereutes donnezani* † | E | Depéret, 1890 |
| **Nyctereutes megamastoides* † | E | Pomel, 1842 |

| | | |
|---|---|---|
| **Speothos pacivorus* † | B | Lund, 1839 |
